# Insights into assembly of constitutive filament-forming proteins of novel fold through domain dissection and EGFP insertion-based screen: A case study of *Spiroplasma* Fibril

**DOI:** 10.1101/2023.10.26.564286

**Authors:** Mrinmayee Bapat, Shrikant Harne, Rajnandani Kashyap, Pananghat Gayathri

## Abstract

*Spiroplasma* is a cell-wall less, helical bacterium possessing a cytoskeletal ribbon consisting Fibril and 5 MreB paralogs. In absence of any information regarding the filament interface of Fibril, a cytoskeleton protein of novel fold, multiple short constructs of Fibril were designed to identify minimal domains required for polymerization. Purification trials of these short constructs resulted in insoluble protein products. In a parallel approach, we incorporated EGFP at the loop regions aimed to hinder polymerization and obtain non-polymerising constructs. We were able to identify probable interfilament/intrafilament interface residues based on the fluorescence screen. We show that EGFP insertions at many of the positions are tolerated as Fibril filaments were readily observed for these constructs in transmission electron microscopy. Both the approaches suggest that Fibril requires N and C-terminal domains for its polymerization. Earlier studies have shown that Fibril^WT^ interacts with MreB5. Our Fibril-EGFP constructs and MreB5 showed interaction similar to the wild-type. Co-sedimentation assay and visualization of Fibril-EGFP proteins proves that the fluorescent constructs are folded and functional and possess structural and biochemical properties similar to the wild type protein.

## Introduction

Actin, tubulin and intermediate filaments are the three major types of proteins that constitute the eukaryotic cytoskeleton (Mitchison, 1995). Typically, actin, tubulin and their prokaryotic homologs polymerize upon nucleotide-binding (ATP and GTP respectively) and depolymerize upon nucleotide hydrolysis (Amos et al., 2004). Conversely intermediate filaments and some prokaryotic cytoskeletal proteins form constitutive/stable nucleotide-independent polymers (Lin and Thanbichler, 2013). In addition to the relevance for function, the control over polymerization and de-polymerization by nucleotides or ligands is useful for purification and characterization of these proteins (Arai et al., 1975; Nonomura et al., 1975). The knowledge about fold and probable polymerization interface based on homology to the eukaryotic proteins is helpful for designing non-polymerizing mutants. These help in designing experimental strategies for structural studies of monomeric conformations, and hence in understanding assembly. Thus actin, tubulin and their homologs are well studied among cytoskeletal proteins. In contrast, the constitutive nature of nucleotide-independent filaments makes them challenging to purify and characterize *in vitro* with respect to understanding their mechanism and dynamics of polymerization assembly. Crescentin, bactofilin, DivIVA, ESCRT-III, CrvA, etc. are some of the proteins found in prokaryotes that form constitutive filaments (Kühn et al., 2010; Wickstead and Gull, 2011).

Many of the constitutive, nucleotide-independent filament forming proteins cannot be depolymerized by known methods and the filaments are not amenable for crystallization. Therefore, they are not suitable for characterization using X-ray crystallography. These structures are challenging to determine using cryo-electron microscopy (cryoEM) when isolation from natural sources is difficult. Latest advances in structure prediction such as AlphaFold also finds it challenging to predict these structures accurately because these exist as higher order oligomers. Overexpression strategies lead to bundles of constitutive filaments that are challenging to purify as well-separated individual filaments, suitable for 3D reconstruction. Structural information on non-polar filaments of bactofilin and crescentin have been obtained recently using cryoEM through modification strategies using nanobody derivatives that led to the formation of non-bundled individual filaments (Liu et al., 2023; Shi et al., 2015).

### Spiroplasma

Fibril is a bacterial cytoskeletal protein, which is hypothesized to be a decisive factor for conferring the helical shape to the cell wall less bacterium. It is a protein of unknown fold, which forms part of a central cytoskeletal ribbon within the cell. Evidence from cryotomography showed that the cytoskeletal ribbon consists of MreB, the bacterial actin, in addition to Fibril (Kürner, et al., 2005). Conformational changes of the Fibril filament have been proposed to play a role in driving the change in handedness during kinking motility in *Spiroplasma* (Sasajima et al., 2022; Shaevitz et al., 2005). It was suggested that Fibril might be responsible for the cork-screw movement of the helical organism because of the force generated by attaching to the cell membrane (Cohen-Krausz et al., 2011). Recent studies show that the helical pitch of the Fibril ribbon closely matches the helicity of the cell body of *Spiroplasma eriocheris* (Sasajima et al., 2022). It has been demonstrated that combination of MreB paralogs along with Fibril was sufficient to cause helicity and motility in synthetic cells (Kiyama et al., 2022; Lartigue et al., 2022). There is also a possibility that Fibril may not have a role in helicity determination in some *Spiroplasma*, where the presence of Fibril gene has been compensated by additional MreBs, for example in *S. sabaudiense* and *S. helicoides* (Abalain-Colloc, et al., 1987; Harne et al., 2020; Shen et al., 2016).

Fibril is a 59 kilodalton protein in which nearly half of the sequence (residues 1–234) at the N-terminus shares homology with MTAN (5’-methylthioadenosine nucleosidase) (Cohen-Krausz et al., 2011). The C-terminus of the protein (235-512) shares no homology and was of unidentified fold. Recent structure determination of Fibril filament from *Spiroplasma eriocheris* will help in fold identification of the C-terminal domain (Sasajima et al., 2023). Since Fibril is a cytoskeletal protein of novel fold, it was challenging to identify the residues at the polymerization interface in the absence of structural information. Interestingly, even latest advances in structure prediction such as AlphaFold did not predict the structure of fibril monomers reliably, especially for the C-terminal domain.

We had recently reported the purification of *Spiroplasma citri* Fibril, heterologously expressed in *E. coli* (Harne and Gayathri, 2022). With a goal to study the filament assembly of fibril, we aimed at obtaining well folded, non-polymerizing constructs of fibril as well as a fluorescently-tagged construct that assembles into a functional filament. In absence of any information about domain boundaries and polymerization interface, we planned to investigate two approaches to identify different domains of Fibril and characterize them using structural and functional approaches. The first approach was based on the domain-wise dissection of the protein into shorter constructs based on its homology with 5’-methylthioadenosine nucleosidase (MTAN; PDB ID 1JYS). In another parallel approach, we introduced EGFP at different sites of the protein to inhibit polymerization and obtain a well folded non-polymerizing construct. In addition to providing a non-polymerizing construct and interface information, this strategy also will help in identifying positions at which fluorescent tag insertion is tolerated within the fold of the protein in a polymerized state. This is a crucial aspect for future studies in visualizing Fibril filaments within the cell.

Monomeric EGFP was used as a fluorescent tag to make sure that the oligomerization of EGFP will not lead to artefacts (Zacharias et al, 2002). The fluorescence intensity is indicative of the folding of the protein of interest (Gregoire et al., 2012; Waldo et al., 1999). If the insertions interfere with the polymerization of the protein, we expect to get a non-polymerizing construct, the fluorescence intensity being a reporter of the amount of folded protein. GFP based screens for *E. coli* membrane proteins (MP) have shown that the whole-cell fluorescence is a useful indicator for the overexpression of MP-GFP fusions (Drew et al., 2005; Geertsma et al., 2008).

Our results suggest that fibril is an exceptionally stable filament assembly. Domain-wise dissection of fibril did not result in folded protein, suggesting that the full-length protein and polymerization are probably essential for the stability of the protein. Fibril is a protein which possesses a polymerization (intrafilament interface), a probable membrane interaction surface and the interfilament interfaces with other Fibril filaments and MreB. Disruption of any of these interfaces will affect the function of fibril within *Spiroplasma*. Hence, a EGFP insertion-based screen is a valuable approach to identify the best position of insertion of EGFP for in vivo studies without disruption of the filaments and functional interactions. This will be useful for shortlisting positions for EGFP insertion for a functional complementation of fluorescently tagged fibril monomers.

## Material and methods

### Sequence analysis

DNA and protein sequences of *S. citri* GII-3 *fibril* (Sequence ID: AM285313.1) were used as query for identification of closely related proteins using BLAST (https://blast.ncbi.nlm.nih.gov/Blast.cgi?PAGE=Proteins). Secondary structure of fibril was predicted using psipred (http://bioinf.cs.ucl.ac.uk/psipred/) and used as the basis for preparation of short constructs as well as for deciding sites of insertion of EGFP.

### Cloning

The *fibril* construct of *S. citri* reported in (Harne and Gayathri, 2022) was used as the template for preparation of short constructs as well as *egfp*-tagged constructs in pHis17 vector (Addgene plasmid #78201) by restriction-free (RF) cloning method (van den Ent and Lowe, 2006) using appropriate primers (Supplementary table 1). All the clones were sequenced using T7 primers to confirm sequences.

### Expression check of *fibril* constructs and *fibril-egfp* constructs

All the proteins were overexpressed using *E. coli* BL21 (AI) cells (Invitrogen) by growing the transformants in Luria Bertani (LB) broth supplemented with ampicillin (100 µg/mL final concentration) at 37 °C under shaking conditions until OD_600_ reached 0.6. The conversion of optical density of the bacterial cultures to cell density was done using Agilent *E. coli* bio calculator (https://www.agilent.com/store/biocalculators/calcODBacterial.jsp). All the cultures were then induced with sterile L-arabinose (final concentration 0.2 %) to initiate gene expression. Post-induction full-length *fibril* and the short constructs were grown at 25 °C for 6 hours while the EGFP-tagged full-length *fibril* constructs were grown at 18 °C for 12 hours. 10 mL aliquots of growing culture (induced and uninduced) were pelleted separately and used for expression check. The cultures expressing protein of interest was pelleted by spinning at 21,000 g at 4 ⁰C and stored at −80 ⁰C until further use.

### Purification of proteins

#### Purification of Fibril-EGFP using density gradient

EGFP-tagged full-length fibril constructs were purified using the urografin density gradient (Harne et al., 2020; Harne and Gayathri, 2022). Cells expressing desired protein were lysed by sonication and spun at 4629 g at 4 ⁰C for 10 minutes to get rid of the cell debris as pellet. The clarified lysate was further spun at 159,000 g at 4 ⁰C for 30 minutes to obtain Fibril polymers in the pellet. The pellet was re-suspended in T_10_E_10_ buffer (10 mM Tris pH 8.0 and 10 mM EDTA) and stirred at 4 ⁰C about 20 hours. The stirred solution was separated on urografin gradient and urografin was removed from the protein by cycles of pelleting and re-suspension in 50 mM Tris to obtain enriched Fibril.

#### Purification of Fibril Δ(305-512)-His_6_

A 400 mL culture pellet of *E. coli* cells expressing His_6_-Fibril Δ(305-512) construct was re-suspended in 100 mL lysis buffer containing 10 mM Tris pH 7.6, 1 % Triton X-100 and 0.5 % LSS and lysed by sonication. The lysate was spun at 7000 g at 4 ⁰C for 15 minutes to separate cell debris. In order to separate Fibril from other *E. coli* proteins, the supernatant was transferred to a fresh tube and spun at 159,000 g at 4 ⁰C for 30 minutes. The pellet was re-suspended in 1.9 mL buffer containing T_10_E_1_; 10 mM Tris pH 7.6 and 1 mM EDTA and 100 μL SDS solution (20 % w/v) for solubilizing the pellet. The solution was then spun at 21,000 g at 25 ⁰C for 15 minutes to get rid of insoluble matter. A linear glycerol gradient was prepared consisting of 17 mL each of 10 % and 50 % glycerol in T_10_E_10_ buffer using gradient mixer.

1 mL of the supernatant obtained in the previous step was loaded onto the pre-cast glycerol gradient. The gradients were then spun at 79,500 g at 4 ⁰C for 2 hours to achieve protein separation. The gradient was then fractionated using peristaltic pump assisted by a glass capillary. The fractions were sequentially labeled from bottom of the tube to the topmost layer. Protein from all the fractions was visualized on SDS-PAGE gels and fractions containing protein of interest were pooled. The protein fractions were dialysed against 2 L of T_10_E_10_ buffer for removing glycerol.

#### Purification of Fibril Δ(273-512)-His_6_ with SDS

400 ml culture pellet of *E. coli* cells expressing truncated Fibril construct was thawed on ice. The pellet was re-suspended in 50 ml of lysis buffer containing 10 mM Tris pH 7.6, 1 % Triton X-100 and 0.5 % LSS. Cells were lysed by sonication for 18 minutes with cycles of 1 second ON, 3 seconds OFF with 60 % amplitude. EDTA was added to the lysate at final concentration of 0.5 mM to prevent non-specific binding of proteins to Ni-NTA matrix. The lysate was spun initially at 14,600 g at 4 ⁰C for 15 minutes to remove cell debris. The supernatant fraction containing protein of interest was further spun at 72,700 g at 4 ⁰C for a duration of 30 minutes to separate Fibril filaments as pellet from other non-polymerized *E.coli* proteins. The pellet containing Fibril was re-suspended in 1.9 mL of T_10_E_1_ buffer. 100 µL SDS (20 % w/v) solution was added along with the resuspension buffer for solubilization of the pellet. The solution was spun at 21,000 g at 25 ⁰C for 15 minutes to get rid of insoluble particulate matter. The supernatant was further spun at 159,000 g at 25 ⁰C for 15 minutes to separate aggregates/smaller clumps as pellet. The supernatant containing Fibril was dialyzed three times against 2 litres of distilled water for 1 hour to remove SDS, EDTA and other salts. The dialyzed protein was transferred to a fresh tube and Tris pH 7.6 and NaCl were added at the final concentrations of 50 mM and 200 mM respectively. The protein was then mixed with Ni-NTA resins pre-equilibrated with binding buffer containing 50 mM Tris pH 7.6 and 200 mM NaCl and incubated by mixing on rotomix (Tarsons) at room temperature (25 ⁰C) for 3 hours. In case of low affinity for binding to the resins, the incubation allows histidine tag of the proteins to bind to Ni-NTA resins more effectively. Following incubation, the mixture was loaded into a syringe column (10 ml) and eluted using Elution buffer containing imidazole in binding buffer. Elution buffer consisting of (25 mM, 50 mM, 100 mM, 250 mM and 500 mM) imidazole was passed through the resin in steps and each fraction was collected separately. 10 µL from each of these fractions were added to 10 µL 2X SDS-PAGE loading and incubated at 95 ⁰C for 10 minutes with shaking at 300 revolution per minute (rpm). Samples were briefly spun and 10 µL loaded onto the 12 % SDS-PAGE gel along with a protein marker.

Fractions containing purified protein were pooled and dialyzed thrice against 2 litre distilled water for 1 hour to remove salts and imidazole. The dialyzed protein was concentrated using centricons with cut off range of 3 kDa. The concentrated protein was aliquoted as 20 µL PCR tubes and flash frozen in liquid nitrogen. the protein was stored at −80 ⁰C until further use.

#### Purification of Fibril Δ(273-512)-His_6_ without SDS

A 2-litre culture pellet of *E. coli* cells expressing Fibril Δ(273-512)-His_6_ construct was thawed on ice. The pellet was re-suspended in 200 mL lysis buffer containing 50 mM Tris pH 8.0, 200 mM NaCl and 0.5 % LSS. Cells were lysed by sonication for 48 minutes with cycles of 1 second ON, 3 seconds OFF with 60% amplitude. The lysate was spun at 46,500 g at 4 ⁰C for 15 minutes to remove cell debris. The supernatant containing Fibril Δ(273-512)-His_6_ was loaded into a pre-packed 5 mL Ni-NTA column (HisTrap; GE Healthcare) equilibrated with binding buffer, 50 mM Tris pH 8.0 and 200 mM NaCl using Äkta system in a cold cabinet (4 ⁰C). The unbound protein was collected as flowthrough. 40 column volumes (200 mL) of binding buffer were passed through the column to remove any unbound proteins and collected as wash. Bound protein was eluted from column using increasing concentrations of elution buffer 50 mM Tris pH 8.0, 200 mM NaCl and 500 mM imidazole. 8 column volumes each were collected with 10 mM and 20 mM imidazole containing buffer. 6 column volumes were collected for buffer containing 50, 100, 250 and 500 mM imidazole each. The eluted protein was collected as 5 mL fractions. 10 µL sample was taken at each step and mixed with equal volume of 2X SDS-PAGE loading dye. The samples were heated at 99 ⁰C for 10 minutes at 350 rpm and spun briefly before loading on 12 % SDS-PAGE gel. Fractions containing protein of interest were pooled and spun at 39,139 g at 4 ⁰C for 30 minutes for removal of any precipitation. The supernatant protein containing protein of interest was transferred into a dialysis membrane with a 3 kDa cut-off. The dialysis was performed against 2 litres of a low salt buffer A50 (50 mM Tris pH 7.6 and 50 mM NaCl) for 1 hour to remove salts and imidazole. The dialyzed protein was again spun at 39,139 g at 4 ⁰C for 30 minutes to get rid of precipitated proteins. The soluble protein was loaded onto a 1.7 mL cation-exchanger column (MonoS; GE Healthcare) pre-equilibrated with buffer A50 (50 mM Tris pH 7.6, 50 mM NaCl). The unbound protein was collected as flowthrough. The bound protein was eluted by passing 20 column volumes (34 mL) of linear gradient between 0 % and 100 % of elution buffer (50 mM Tris pH 7.6 and 500 mM NaCl) against binding buffer (50 mM Tris pH 7.6 and 50 mM NaCl). 1 mL fractions were collected during elution. Samples from each step were checked on 12 % SDS-PAGE gel. The flowthrough from MonoS column was spun at 39,139 g at 4 ⁰C for 30 minutes to remove precipitated proteins. Since the protein did not bind to MonoS column, it was passed through an anion exchange column (MonoQ; GE Healthcare) column to check if can bind to the latter. The supernatant was loaded onto MonoQ pre-equilibrated with buffer A50 (50 mM Tris pH 7.6 and 50 mM NaCl. The unbound proteins were collected as flowthrough and the bound proteins were eluted using the buffers and protocol described for MonoS purification. The fractions were visualized on 12 % SDS-PAGE gels. The fractions containing protein of interest were pooled and concentrated and stored at −80 ⁰C.

### Pelleting assay of short constructs of Fibril

To verify if a protein construct forms polymers or not, we performed centrifugation-based pelleting/pelleting assay. Filamentous proteins have been shown to pellet down upon spinning at force > 80,000 g. However, the non-polymeric protein will be obtained in supernatant and is often used as a purification method for polymeric proteins (Harne and Gayathri, 2022; Mizuno, 1992; Woodring et al., 2001). Based on these observations, oligomerization state of short constructs was also monitored using pelleting assay.

For pelleting assay, 400-mL culture pellet of cells expressing protein of interest were lysed in 50 mL lysis buffer (10 mM Tris pH 7.6, 1 % Triton X-100, and 0.5 % LSS) by sonication. EDTA was added in the lysate at a final concentration of 1 mM to depolymerize nucleotide-dependent polymers of proteins. The lysate was initially spun at 14,636 g at 4 ⁰C for 15 minutes to remove cell debris as pellet. The soluble fraction was spun at 46,500 g at 4 ⁰C for 30 minutes to pellet down Fibril filaments. Pellet was re-suspended in 2 mL of T_10_E_1_ (10 mM Tris pH 7.6 and 1 mM EDTA buffer and spun at 21,150 g at 25 ⁰C for 15 minutes to pellet down insoluble aggregates. The soluble fractions were transferred to a fresh tube and spun at 159,000 g for 15 minutes at 4 ⁰C to pellet down filamentous proteins. The supernatant was transferred to a fresh tube while the pellet was re-suspended in 40 μL distilled water. 10 μL sample was taken at each step during the purification and checked on a 12 % SDS-PAGE gel.

### Estimation of whole cell fluorescence

All the EGFP-tagged Fibril constructs were transformed in BL21 (AI) cells and incubated at 37 °C overnight. Cells were further grown in LB media supplemented with 100 µg/mL ampicillin. For calculating cell density from optical density of the cultures, OD_600_ calculator from Agilent was used (https://www.agilent.com/store/biocalculators/calcODBacterial.jsp). When the cultures attained optical density (cell density: −4.8*10^8^) OD_600_ - 0.6, they were induced with sterile L-arabinose at a final concentration of 0.2 % w/v and further incubated at 18 °C for 12 hours. The cultures were normalized to a cell density of 4.8 × 10^8^ before induction. 1.5 mL of culture was pelleted down and washed twice with 1X phosphate buffered saline (PBS, 137 mM NaCl, 2.7 mM KCl, 10 mM Na_2_HPO_4_ and 1.8 mM KH_2_PO_4_; pH 7.4) to remove traces of LB media and final pellet was re-suspended in 300 µL PBS (1X) each. All the cultures were transferred into a 96 well plate and OD_600_ measured to re-confirm the equal cell density for all the cultures. Same cultures were then transferred to a black plate for measuring whole cell fluorescence intensity. The excitation and emission wavelengths for measuring the fluorescence were 488 nm and 509 nm respectively. The fluorescence intensity was plotted against normalized cell density of 2 × 10^9^ (OD_600_ – 2.5) for different EGFP-tagged Fibril constructs.

### Confocal Microscopy

The EGFP-tagged Fibril constructs were grown as described above. The cell densities (as estimated by measuring OD_600_ of cultures were diluted to achieve OD_600_ = 2 (cell density = 1.6 × 10^9^). 1 mL of each culture was pelleted and cells washed twice with 100 µL PBS (1X). The pellets were re-suspended in 100 µL paraformaldehyde (4 %) and incubated at room temperature for 30 minutes. Cells were then spun at 15871 g for 1 minute and pellets were washed twice with 1X PBS to remove excess of paraformaldehyde and washed cells re-suspended in 100 µL 1X PBS. 2 µL of this solution was used to prepare slides and kept for drying at room temperature. 5 µL of 80 % glycerol was added above the smear as a mountant and sealed with coverslip. The slides were then used for visualization using a confocal microscope (Leica SP8) with Argon laser (20 %) with the parameters as gain 874.9, offset 0.1 %, pinhole 1 AU, line average 2. The settings for microscope were kept constant for visualizing all the constructs.

The fluorescence intensities and cell sizes were quantified using ImageJ (Schneider et al., 2012). Individual cells were selected using the DIC filter as the cell boundaries were clearly seen. Mean grey value and length were calculated for approximately 50 cells for each construct and plotted against different constructs.

### *In vitro* fluorescent intensity estimation of Fibril-EGFP protein

Fibril-EGFP proteins purified by urografin density gradient were fractionated in equal volumes and added in a Corning 96-well plate. 300 μL fractions were aliquoted starting from the top of the gradient and added in the plate. Scattering was measured at 600 nm. All the fractions were then transferred to Costar 96-well black plate. Fluorescence readings were measured at E_ex_ - 488 nm and E_em_ - 509 nm. All readings were measured in EnSight multimode plate reader (Perkin Elmer).

### Sample preparation for CD spectroscopy

Samples for Fibril 27 EGFP, Fibril 183 EGFP and Fibril 493 EGFP were prepared for CD spectroscopy. The proteins were treated with DNAse at 37°C for 1 hour. The protein was spun at 21000 g for 10 minutes and supernatant was discarded. Proteins were then resuspended in equal volumes of MilliQ water. 9.5 μL of the protein was added to 0.5 μL 1% SDS. Concentration of protein was estimated using Nanodrop. Solution of 9.5 μl MilliQ water and 1% SDS was used as blank. 500 μL of 5 μM of each protein were transferred to fresh tubes containing 1% SDS and MilliQ water. The spectra were recorded in JASCO J-815 CD spectrophotometer using 1 mm cuvette, scan speed of 50 nm/min, digital integration time of 2 s and a bandwidth of 1 nm. Spectra scan was done from 200 nm to 250 nm. 20 spectra were averaged for each protein.

### Sample preparation and visualization of Fibril filaments using Transmission Electron Microscopy (TEM)

2 % (w/v) uranyl acetate (Sigma-Aldrich) was prepared by dissolving 200 mg powder in 10 mL sterile distilled water. The solution was filtered using 0.22 μm filter and then distributed as 500 μL aliquots into 1.5 mL tubes. All the uranyl acetate containing tubes were covered with aluminum foil and stored at 4 ⁰C until use. For sample preparation, solution of desired concentration (0.5 % w/v) was prepared by diluting the stain solution with sterile water.

For short constructs, Carbon-Formvar coated copper grids (Ted-Pella, Inc) were glow-discharged in plasma cleaner (Quorum technologies) just before use. 10 µL of sterile water was loaded on the grids to make them hydrophilic. Varying volumes (2 - 5 μL) of purified protein was added to the water present on the grid and allowed to stand at room temperature for at least 30 - 180 seconds to facilitate settling down of the filaments. For Fibril-EGFP constructs, carbon-formvar coated copper grids (200 mesh) were used without glow discharge. 10 μL of protein was loaded on the grid for 3 minutes. Excess buffer was back-blotted using Whatman filter paper. The grid surface was washed using 10 μL of MilliQ water and back blotted. 4 μL of stain (0.5 % w/v uranyl acetate) was applied to the grid and absorbed from bottom using blotting paper after 30 seconds so as to prevent absorption of the stain by protein. The grids were allowed to air dry at room temperature in a dust-free environment and used for observation using TEM.

### Pelleting assay of Fibril-EGFP and MreB5^WT^

Fibril-EGFP and MreB5^WT^ interaction were checked with three of the fluorescent constructs, namely, Fibril 27 EGFP, Fibril 304 EGFP and Fibril 493 EGFP. Varying concentrations of Fibril-EGFP were used whereas MreB5^WT^ concentration was kept constant. For the assay, 10 μM of MreB5^WT^ was used. MreB5^WT^ was purified as reported in (Harne et al., 2020; Pande et al., 2022) and maintained in a final buffer of 300 mM KCl, 50 mM Tris pH 8.0. The range of Fibril-EGFP concentrations used were 2.5 μM, 5 μM, 10 μM, 20 μM, 40 μM and 50 μM. MreB5^WT^ concentration was estimated by checking absorption at 280 nm. Fibril-EGFP concentration estimation was done by dissolving 9.5 μL of purified protein in 1 % SDS. The concentration was estimated by checking the absorption at 280 nm. MilliQ with 1 % SDS was used as a blank for concentration estimation. The two proteins were added together in a buffer containing in 300 mM KCl, 50 mM Tris pH 8.0. 10 μM MreB5^WT^ and 2.5 μM to 50 μM Fibril-EGFP were kept as protein controls. The reactions were incubated at 25 ⁰C for 30 minutes. Post incubation, the reactions were spun at 100000 g for 15 minutes at 4 ⁰C. The samples were spun in TLA 120.2 rotor in Optima Max-XP ultracentrifuge (Beckman Coulter). Supernatant was transferred to a fresh tube and pellet was washed with 100 μL of 300 mM KCl, 50 mM Tris pH 8.0. Pellet was resuspended in 50 μL buffer. 10 μL of each supernatant and pellet were added to Coomassie brilliant blue dye and heated at 99 ⁰C for 10 minutes. The samples were loaded on a 10 % SDS PAGE gel.

## Results

### Domain-wise dissection of Fibril affects protein stability and solubility

The shorter constructs of Fibril were designed on the basis of secondary structure prediction using PsiPred (Figure 1A). The boundaries of N-terminal and C-terminal domains were identified based on the sequence alignment between Fibril and MTAN (PDB ID: 1JYS) (Figure 1A and S1). For designing short constructs of Fibril, loop regions, highlighted in grey, were selected as domain boundaries (Figure 1A and 1B). Similar to full length fibril, short constructs of Fibril were seen to pellet at 46500 g (Figure 1C). Further purification of the protein obtained in pellet were tried using density gradient centrifugation and affinity chromatography. The protein purity was seen to improve with this technique (Figure S2). However, the proteins purified from short constructs of Fibril were highly unstable and were prone to precipitation, and the results showed batch to batch variation. Fibril shows the presence of an N-terminal and a C-terminal domain. Deletion of a portion of C-terminal domain led to unstable or unfolded proteins. This indicates that polymerization and stability of the protein requires the presence of both the domains of the protein. Fibril might not fold correctly or polymerize when either of the domains are deleted. The domain-wise deletion approach for obtaining a minimal polymerizing construct of fibril was not successful, indicating that the polymerization interface might not be restricted to either N or C terminal domains, but might be composed of multiple interfaces. This has been corroborated by the recent structure of Fibril filament (Sasajima et al., 2023.).

**Figure 1:**
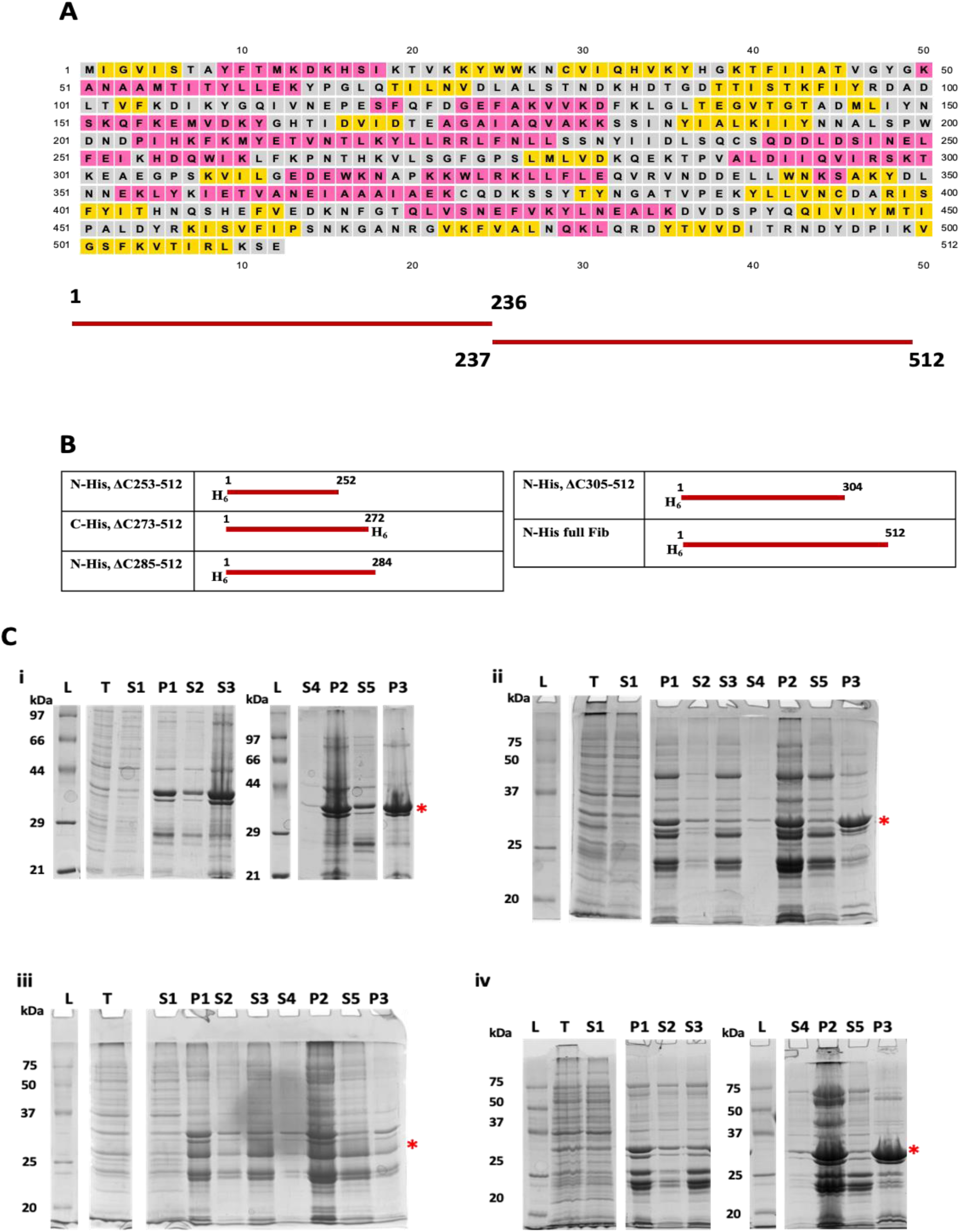
Polymerization/pelleting assay for short constructs of Fibril. **A.** Secondary structure prediction of Fibril was done using Psipred (http://bioinf.cs.ucl.ac.uk/psipred/). N-terminal and C-terminal domains have been highlighted below the secondary structure prediction. **B.** Schematic representation of short constructs **C.** The pelleting assay gels of short constructs His_6_-FibrilΔ(305-512) (**i**; 35kDa His_6_-FibrilΔ(285-512) (**ii**; 33 kDa), FibrilΔ(273-512)-His_6_ (**iii**; 32 kDa) and His_6_-FibrilΔ(253-512) (i**v**; 30 kDa). **L**-Protein marker; **T-** total lysate; **S1-**supernatant upon spinning lysate at 11,000 g; **P1-** Pellet obtained upon spinning protein as in lane 2 at 46,500 g; **S2 & S3**-soluble fraction obtained upon solubilization of the pellet (lane 3) without or with SDS respectively, followed by spinning at 21,000 g; **S4 & P2-** supernatant and pellet respectively after spinning sample from lane **S2** at 159,000 g; **S5 & P3**-supernatant and pellet respectively after spinning sample from lane 5 at 159,000 g.

### Whole cell fluorescence varies across constructs

In order to confirm the observations obtained from short constructs about the domains of Fibril, we took another approach using EGFP as a block at the interface for Fibril polymerization. To gain information about the polymerization interface of Fibril, we prepared multiple fluorescently tagged constructs by insertion of EGFP at various positions. We used the secondary structure prediction tool PsiPred (http://bioinf.cs.ucl.ac.uk/psipred/) to identify the loop regions as sites for EGFP insertion (Figure 2). We observed that all the constructs consistently express EGFP-tagged Fibril as evident from the ∼ 86 kDa bands observed on SDS-PAGE gels (Figure S3).

**Figure 2:**
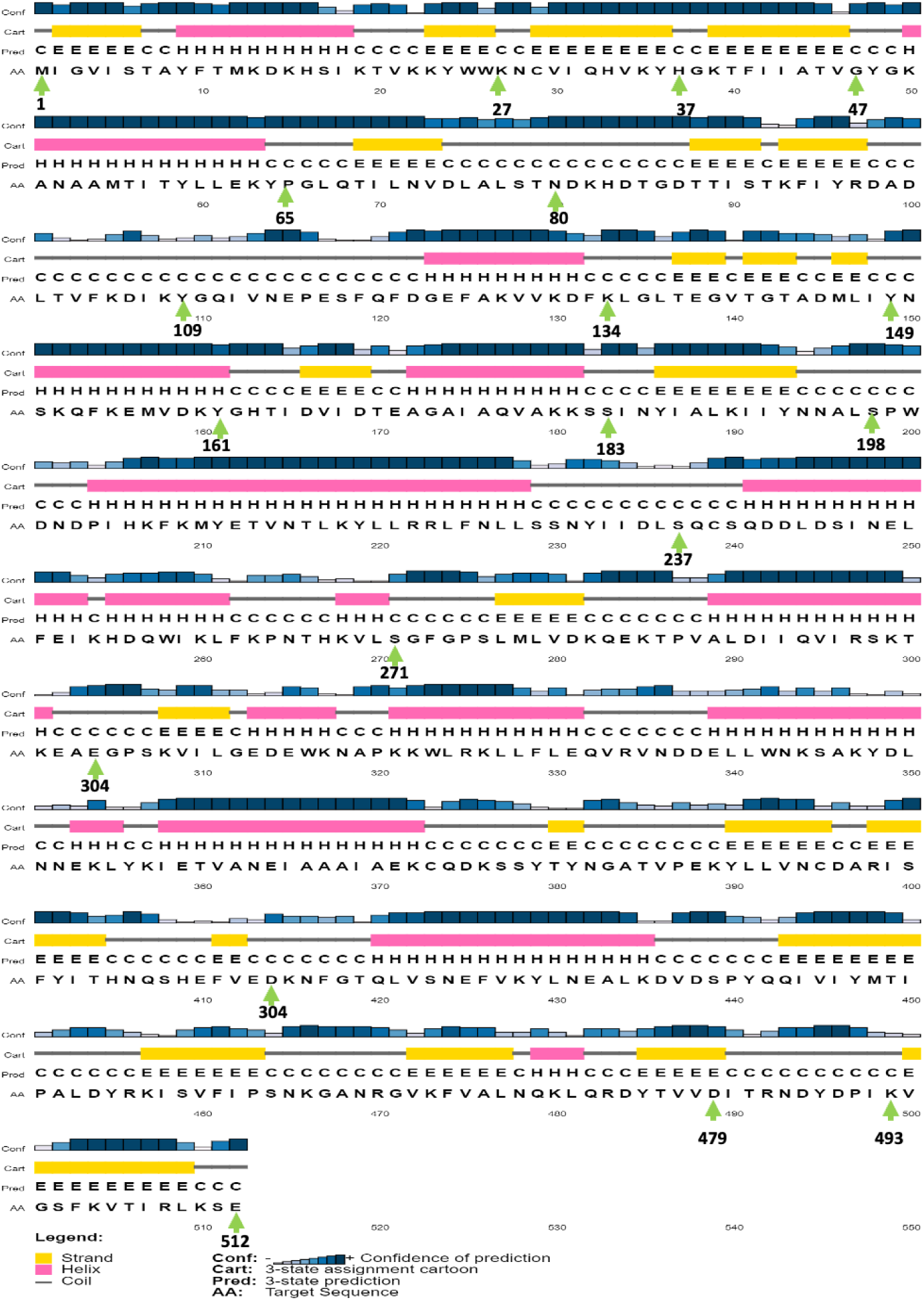
Secondary structure prediction of Fibril. Secondary structure of Fibril was predicted using PSIPRED tool (http://bioinf.cs.ucl.ac.uk/psipred/). Red arrow signifies position of EGFP insertion at the loop regions of Fibril.

We compared the fluorescence intensities across all the Fibril-EGFP constructs as an estimate for the expression of fusion protein. We estimated the bulk whole cell fluorescence of all the EGFP-tagged Fibril constructs after normalizing their cell density. A considerable variation in bulk cellular fluorescence was observed across different constructs (Figure 3A). To confirm if the variation in fluorescence is indeed reflected at individual cell level, we performed confocal microscopy of *E. coli* cells expressing various constructs of EGFP-tagged Fibril (Figure 3B). We observed that unlike EGFP, all these constructs display a non-diffuse localization in the cell, suggesting that they may be forming polymers or aggregates. We further observed that the mean grey value (measured as fluorescence intensity per unit area) of each construct differs and this variation is consistent with the whole cell fluorescence in bulk (Figure 3C). In order to see if the Fibril EGFP constructs has any effect on the cell size of the *E. coli,* the area of the cells selected for measuring fluorescence intensities was calculated (Figure 3D). Consistent with previous results, we saw that there is a variation in fluorescence amongst different constructs in spite of having comparable cell size.

**Figure 3:**
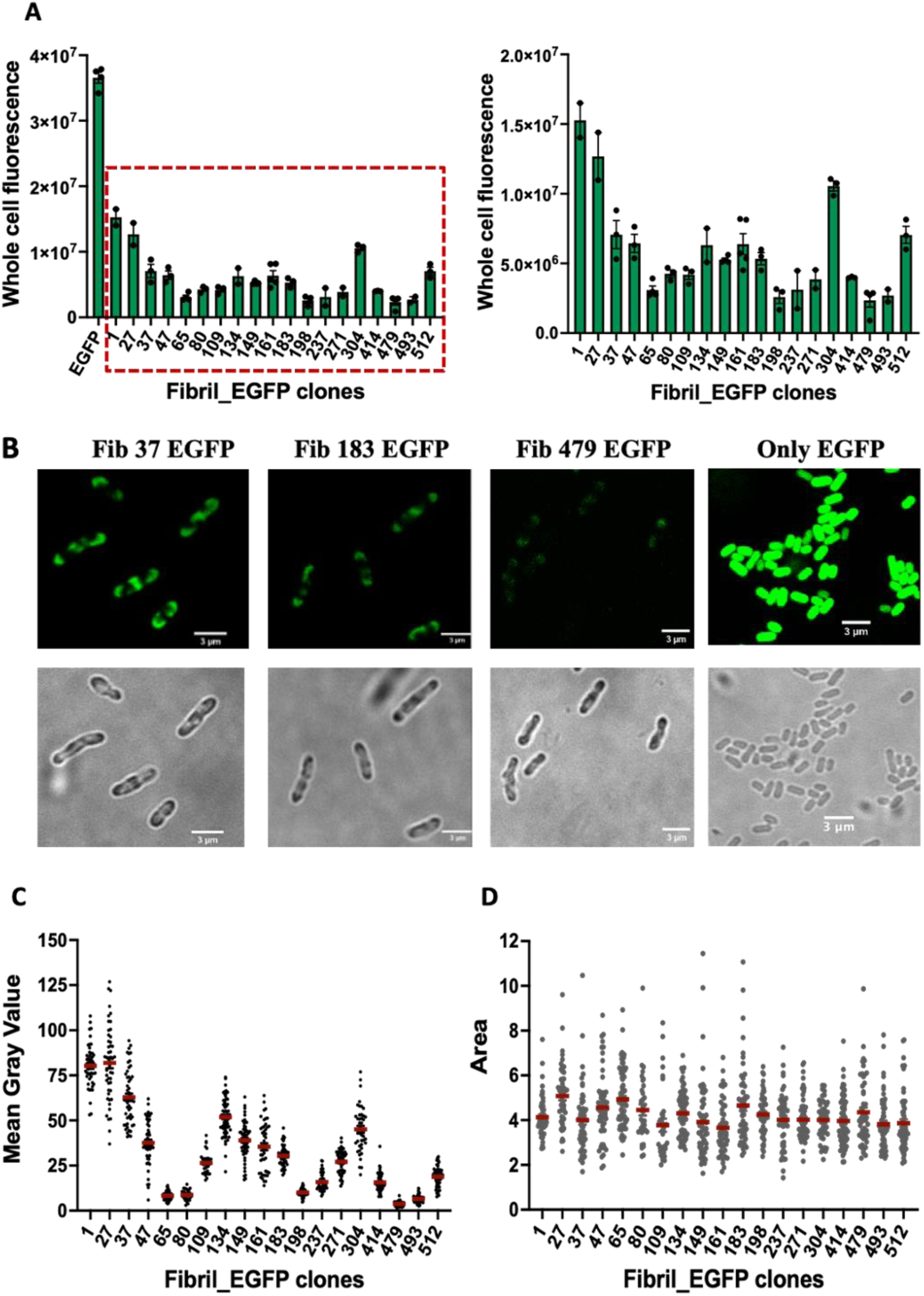
EGFP acts as a folding reporter. **A.** Whole cell fluorescence indicates the folding state of the Fibril-EGFP constructs. The constructs are arranged according to their fluorescence intensities. **B.** Confocal images of 3 Fibril-EGFP constructs. Overexpression of Fibril-EGFP gives a non-diffused pattern of localization whereas only EGFP expression leads to a diffused localization in the cells. **C.** Fluorescent intensities of the individual cells were measured. **D.** Length of the *E.coli* cells were measured to ensure that the size of the cells had not effect on the difference in EGFP fluorescence.

### Protein expression levels of EGFP-tagged constructs of fibril remains similar but the fluorescence varies

The variation in fluorescence could arise from either different levels of expression of the protein constructs or because EGFP tagging of Fibril at certain positions prevents a fraction of protein from folding properly. To verify both these possibilities we checked protein expression, and fluorescent intensities of protein enriched over density gradient ultracentrifugation. We monitored the quantities of protein polymers for each construct by comparing the amount of protein (scattering) and the fluorescence of the fibril filament fractions from density gradient ultracentrifugation. Fibril fractionates into two bands, an upper and a lower band (Figure 4A, B), distinguished by the differences in the proteolytic fragments (Harne and Gayathri, 2022). All the constructs possessed the upper and lower bands in a consistent position of elution suggestive of formation of Fibril polymers of similar densities as the wild type (Figure C) (Harne and Gayathri, 2022). Though the total amount of expressed protein, monitored through scattering intensities, was approximately similar across the constructs, the fluorescence intensities showed a gradation, correlating with the whole cell fluorescence intensities. The protein expression levels across constructs did not show a direct correlation with the amount of fusion protein expressed (Figure 4 D-G). For instance, Fibril-EGFP proteins (eg. Fibril 65 EGFP and Fibril 183 EGFP) showing higher scattering did not show high fluorescence. This suggests that in constructs with relatively low fluorescence, some fraction of the protein molecules must be unfolded aggregates or have the EGFP misfolded.

**Figure 4:**
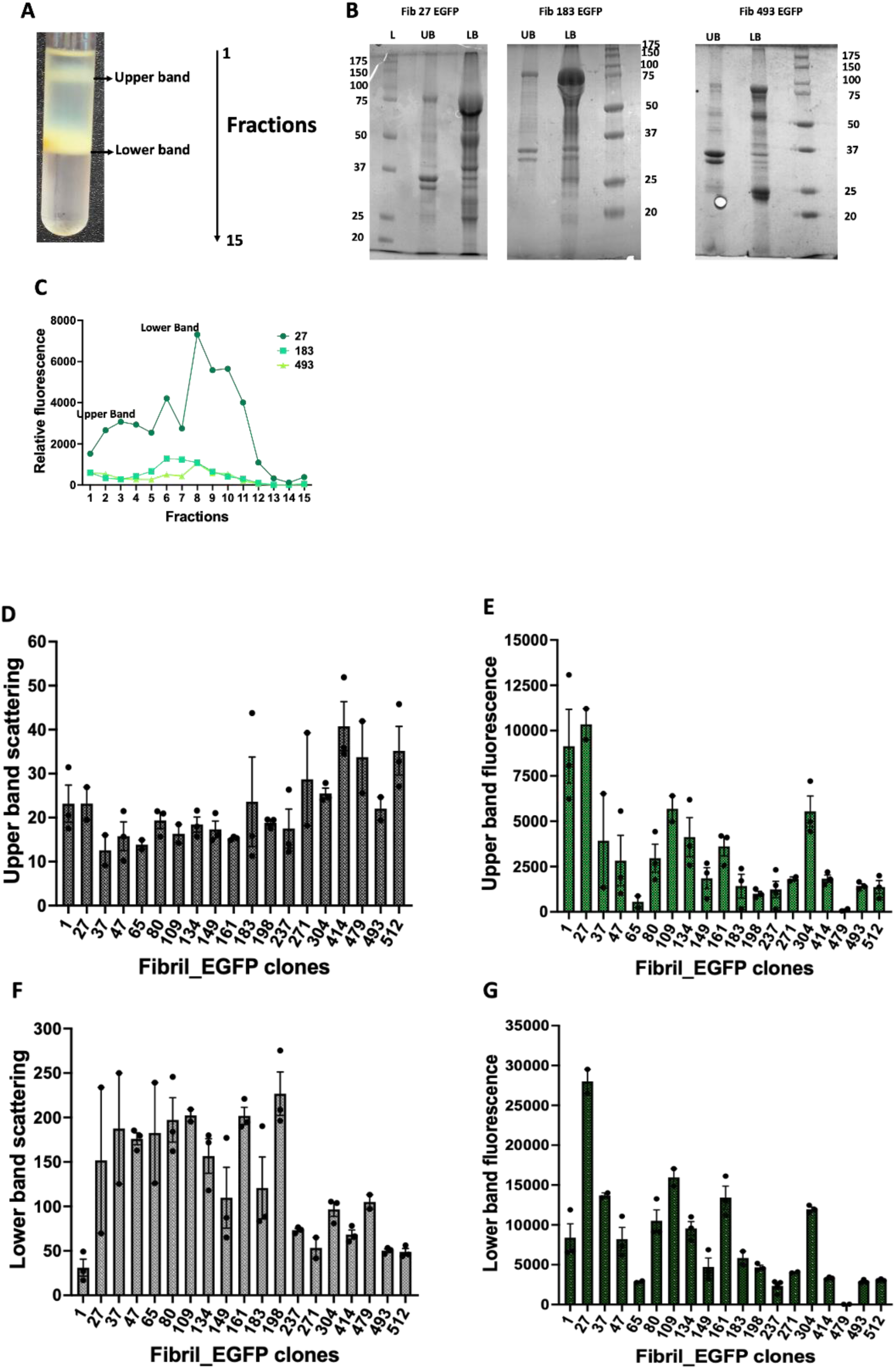
Purification of Fibril-EGFP using urografin density gradient. **A.** Fibril-EGFP purified by urografin density gradient **B.** shows a representative gel of purified Fibril-EGFP constructs using urografin density gradient. **C.** Elution profile of the density gradient for three Fibril-EGFP proteins. The upper and lower band fluorescence correspond to the bands shown in **A.** **D.** Scattering as a measure of total protein amount for upper and lower bands were measured at 600 nm. Fluorescence of the same samples were measured using ***λ***_ex_-488 nm and ***λ***_em_-509 nm.

### CD spectroscopy shows presence of secondary structures similar to Fibril wild type

We relied on CD spectroscopy to monitor the presence of secondary structures compared to the wild type. If Fibril is folded, we expected a signal for α helices as well as β sheets. Three fluorescent proteins, Fibril 27 EGFP, Fibril 183 EGFP and Fibril 493 EGFP at high, mid and low fluorescence levels were chosen as representative constructs. Fibril^WT^ was taken as the control (Figure 5A). Fibril 27 EGFP and Fibril 183 EGFP showed presence of characteristic secondary structures which were similar to Fibril^WT^. The Fibril signature was clearly observed in the CD spectra demonstrating that the Fibril monomers in the filament were folded into the expected globular fold. EGFP gives signal for β sheets alone owing to the 11 β-barrel structure. For Fibril 27 EGFP and Fibril 183 EGFP, the Fibril EGFP combined with EGFP containing 1 % SDS gave spectra which was similar to an additive effect of Fibril^WT^ and EGFP (Figure 5B). However, Fibril 493 EGFP secondary structure signal intensity was seen to be lesser as compared to the other constructs, for the same concentration of protein. This indicates that this particular fusion protein might show presence of a heterogenous population of folded and misfolded protein. This explains a decrease in the CD signal and low fluorescence in whole cell fluorescence and microscopy studies. Equal concentrations of the proteins were used for the analysis, as re-confirmed by analysis of band intensities on a 12 % SDS gel (Figure S4).

**Figure 5:**
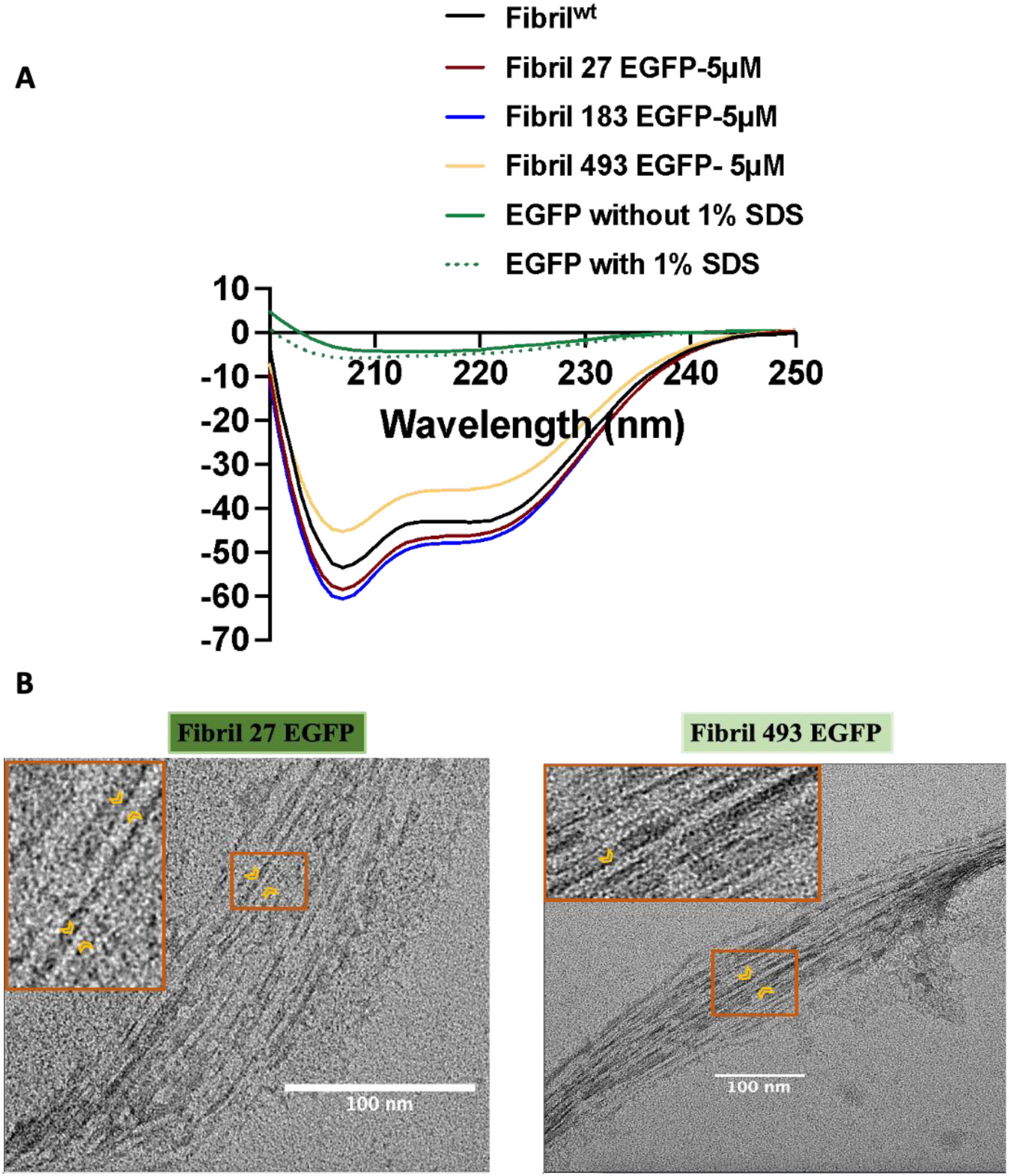
EGFP tagging does not affect secondary structure and filament assembly of Fibril. **A.** Circular dichroism spectra of Fibril 27 EGFP, Fibril 183 EGFP, Fibril 493 EGFP and Fibril^WT^. Three constructs of varying fluorescence levels were selected for checking the effect of EGFP position on Fibril folding. All proteins concentrations were adjusted to 5 µM and were solubilized using 1% SDS. **B.** Negative staining images of Fibril 27 EGFP (highly fluorescent) and Fibril 493 EGFP (low fluorescent). Irrespective of the EGFP position, Fibril were seen to form bundles of filaments. Arrow shows individual filaments. Inset-Zoomed image of protofilament highlighted with the arrows.

Next, we imaged the fluorescently tagged proteins under transmission electron microscopy to confirm that the density gradient fractions of the fusion proteins formed filamentous assemblies and not random aggregates. Fibril 27 EGFP and Fibril 493 EGFP were visualized under transmission electron microscope. Bundles of filaments were seen for both the proteins and the morphology looked similar irrespective of the EGFP tag position. The inset images highlight a single protofilament (Figure 5C). These observations further prove that Fibril readily forms robust filaments irrespective of the position of EGFP insertion. The variation in fluorescence must be result of unfolded EGFP due to steric hindrance of Fibril domains close to the position of EGFP insertion or partially unfolded Fibril.

CD spectroscopy and presence of filaments as seen under transmission electron microscope proves that the purified Fibril proteins are folded and resemble the physical properties earlier observed for wild type protein. The existence of a gradation of fluorescence correlated with the folded secondary structure features confirms that the approach could provide useful insights into the positions where a fluorescent tag might be tolerated more robustly. Thus, these fusion proteins could be used for interaction studies with other cytoskeletal proteins of *Spiroplasma* as an assay for their function.

### Fibril-EGFP interacts with MreB5^WT^

Interaction between Fibril^WT^ and MreB5^WT^ has been shown earlier using co-sedimentation assay (Harne et al., 2020). Fibril^WT^ pellets when spun at 100000 g whereas MreB5^WT^ does not pellet on its own. We performed co-sedimentation assay between Fibril-EGFP for the constructs Fibril 27 EGFP, Fibril 183 EGFP and Fibril 493 EGFP with MreB5^WT^ to see if EGFP tag and its position of insertion has any effect on the interaction between them. Fibril-EGFP protein was obtained in pellet fraction whereas MreB5^WT^ was obtained in supernatant after spinning at 159,000 g, as individual protein components (Figure 6A). The amount of MreB5^WT^ in the pellet fraction increased with an increase in the concentrations of Fibril-EGFP (Figure 6, B-F). These observations were in accordance with the previous result showing interaction between Fibril^WT^ and MreB5^WT^ (Harne et al., 2020). This proves that the Fibril-EGFP proteins are folded and show similar biochemical properties as that of Fibril^WT^. Presence of EGFP at the three different positions did not affect the interaction between the two proteins. This also indicates that EGFP might be away from the interface of the two proteins for these examples.

**Figure 6:**
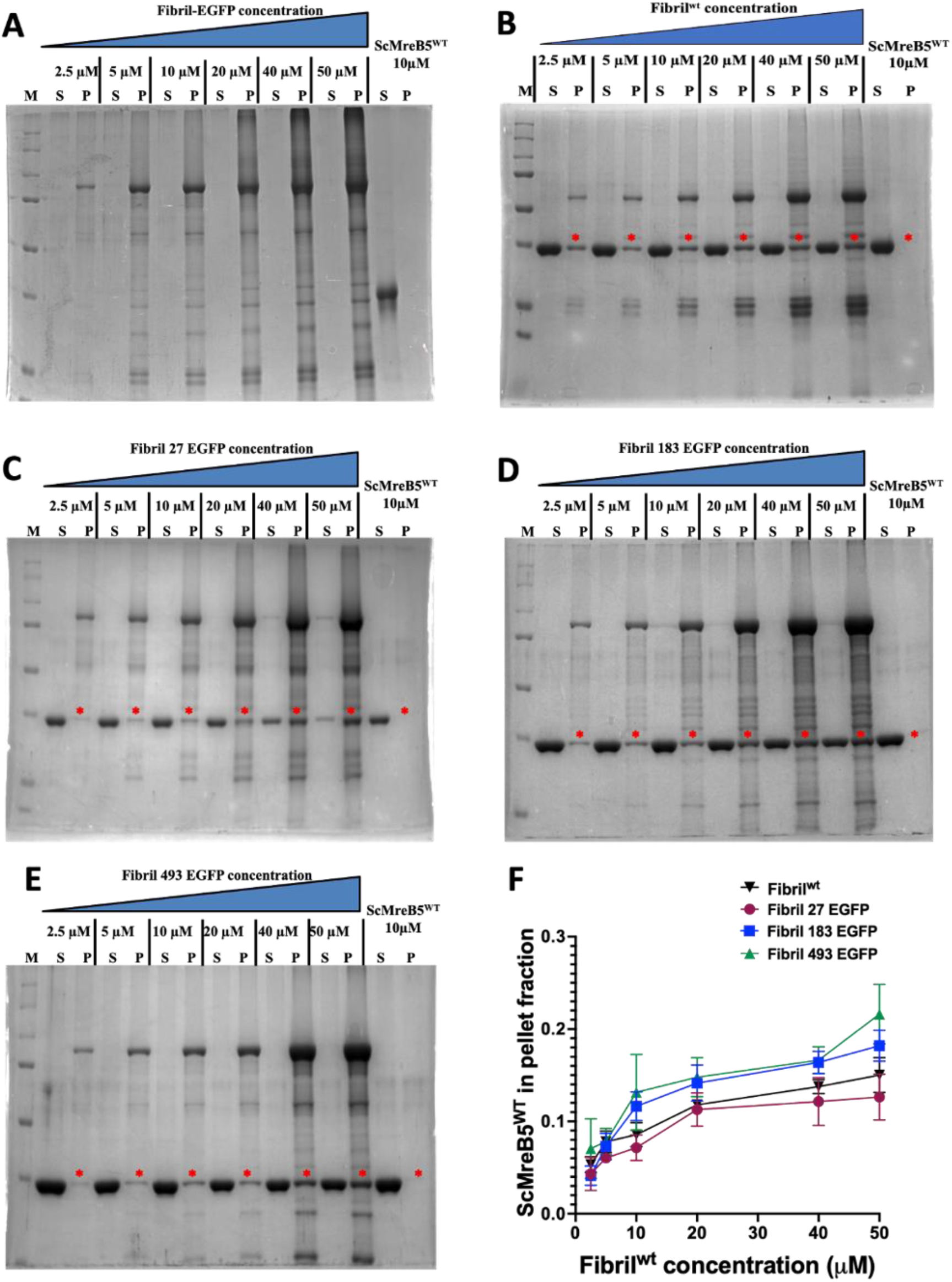
Fibril-EGFP interacts with MreB5^WT^. Fibril^WT^ has been shown to interact with MreB5^WT^ (Harne et al; 2020). Interaction between Fibril-EGFP proteins and MreB5^WT^ was checked to see the effect of EGFP tagging on the interaction of the two proteins. **A – D.** Control gel with Fibril-EGFP in varying concentrations-2.5 µM, 5 µM, 10 µM, 20 µM, 40 µM, 50 µM and MreB^WT^-10 µM. Interaction between Fibril-EGFP and MreB^WT^ were checked by selecting Fibril 27 EGFP (**B**), Fibril 304 EGFP (**C)** and Fibril 493 EGFP (**D**). Three Fibril-EGFP constructs of varying fluorescence were selected for the pelleting assay. Irrespective of the EGFP position, Fibril-EGFP showed interaction with MreB5^WT^ in the absence of any nucleotide. **E.** Quantification of MreB5^WT^ fraction in pellet were plotted against Fibril^WT^ concentration. Increasing interaction of proteins is seen by an increase in pellet fraction. Red star denotes MreB5^WT^ pellet fraction used for quantification.

## Discussion

Whole cell fluorescence and visualization of overexpression of Fibril-EGFP in *E. coli* cells helped us in understanding the effect of insertion of EGFP at different positions of Fibril. This helped us in the identification of residue positions suitable for the addition of EGFP tag. Despite having similar cell densities, these constructs showed varied levels of fluorescence. Amongst all the EGFP insertional positions, Fibril 1, 27, 304 and 37 EGFP were highly fluorescent. It is highly likely that these tag positions accommodate EGFP in its folded state. Fibril 65, 80, 237, 479 and 493 EGFP constructs show the least fluorescence in all our experiments (Figure 7A). Interestingly, these residue positions are not restricted to any particular domain of the protein. AlphaFold model of Fibril dimer shows dimer interface as a part of N-terminal domain alone. C-terminal domain of the model shows a very low pLDDT score (Figure S5) and does not indicate a compactly folded structure. The recent structure also validates that the C-terminal region is not correctly predicted by AlphaFold. Hence, only the N-terminal domain was used for mapping the EGFP positions on the structure. 37, 65, 80, 149, 161, 198 and 237 amino acids seem to be exposed from the model of the N-terminal domain (Figure 7 B and C). There is a possibility that these positions are present between the N and C-terminal domains of the same molecule or at the interface of the two monomers. Due to the steric effect, EGFP has to either move away from the interface space if permissible by the flexible linker or it is likely to misfold. Fibril seem to form highly stable and robust filaments irrespective of the position of EGFP insertion, and calcitrant to unfolding in the presence of the EGFP tags. In some of the constructs, the reduction in fluorescence might be result of loss of EGFP by its proteolytic cleavage leaving the fibril filaments intact. However, all our expression gels showed the presence of a band corresponding to full length protein, which was comparable across the compared constructs.

**Figure 7:**
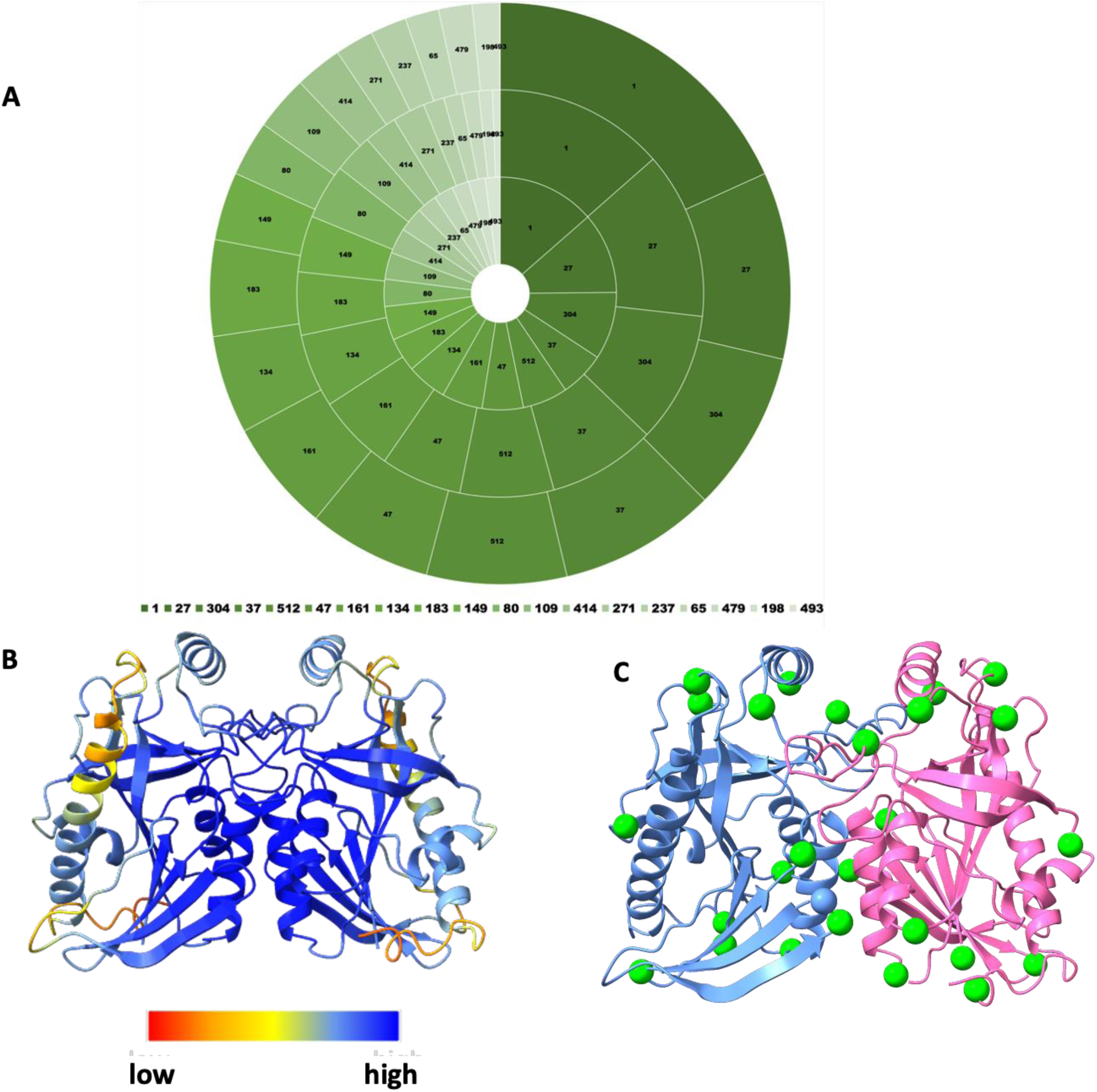
Fibril-EGFP constructs having least fluorescence may have EGFP at the interface. **A.** Pie chart showing profile of fluorescence intensities of different Fibril-EGFP proteins. Outermost circle represents whole cell fluorescence, middle circle represents mean grey value and inner circle represents fluorescence intensity of lower band of purified Fibril. Fluorescence intensities of lower band alone was selected for the comparison as most of the fusion protein is present in the lower band of the gradient. The gradient of green color denotes the intensity of fluorescence in decreasing order in a clockwise manner. **B.** AlphaFold structure of a dimer of Fibril 1-240 amino acid. The structure is color coded according to the pLDDT confidence measure. **C.** EGFP positions were mapped on the AlphaFold model of a dimer of Fibril (1-240 amino acid). EGFP positions at 27, 109, 161 seem to be facing the N-terminal of the adjacent monomer. The helix ranging from amino acid 51-62 also seems to be a part of the interface between two N-terminal domain. However, no EGFP was tagged in this helix region. Fibril 37, 65, 80, 149, 161, 198 and 237 EGFP positions look exposed. These constructs showing low fluorescence may have EGFP close to either C-terminal domain of the same molecule or at the intrafilament/interfilament interface. The C-terminal domain AlphaFold model was not considered for the analysis as it showed a very low pLDDT score.

Since Fibril forms constitutive, nucleotide-independent filaments that cannot be depolymerized without denaturation, obtaining Fibril monomers is challenging. Our results from the domain-wise dissection approach appears to suggest that Fibril is highly unstable in the absence of polymerization. Deletion of a portion from C-terminal domain led to unstable protein constructs. This suggests that the N-terminal domain is not sufficient for polymerization of Fibril. This also suggests that surface residues in Fibril equivalent to MTAN were replaced with hydrophobic residues during the evolution of Fibril by addition of the C-terminal domain.

We used EGFP as a roadblock for polymerization of Fibril. Variation in the fluorescent intensities of Fibril-EGFP fusion proteins might suggest misfolding of the fusion protein or either of the proteins. Transmission electron microscopy of these fluorescently labelled proteins showed that Fibril forms robust filaments irrespective of the EGFP position. This helped us in identifying the position in Fibril where EGFP might be present at the interface of N-terminal domain. These amino acid positions were not specific to any one domain and were present in both N-and C-terminal domain. Based on the observations from short constructs as well as Fibril-EGFP constructs, we suggest that the polymerization interface between two monomers is formed of both N-terminal as well as C-terminal domain. These observations are in agreement with the recent structure of Fibril determined by Cryo-EM (Sasajima et al., 2023).

We observed variation in the fluorescence intensities when Fibril-EGFP were overexpressed in *E.coli* cells. In absence of any information about the structure and a homolog in the database, it is challenging to decide on the position of a fluorescent tag. Many instances have shown that addition of a fluorescent tag results in a non-functional protein (Liu et al., 2023). This also leads to artifacts related to protein localisation in vivo (Jones et al., 2001). Purification of Fibril-EGFP constructs helped us to identify highly fluorescent fusion proteins. Co-sedimentation assay of Fibril-EGFP with MreB5^wt^ showed interaction between the two proteins. This was in accordance with the interactions seen between Fibril^WT^ and MreB5^WT^ (Harne et al., 2020). This further confirms that the fluorescently tagged proteins are well folded and functional. Based on the fluorescent intensities, some of these constructs showing high fluorescence can be used for in vivo localization studies in future.

## Acknowledgements

The authors thank transmission electron microscopy facility at IISER Pune and RCB Faridabad for their constant help with sample imaging. The microscopy facility and staff members of the facility of Biology department of IISER Pune are acknowledged. We thank Suman, Anushka and Biswajit from Prof. Jayant Udgaonkar’s lab for their help with CD spectroscopy.

This project was initiated by the support of funds from Department of Science and Technology (DST) INSPIRE Faculty Fellowship (IFA12/LSBM-52) and Innovative Young Biotechnologist Award (BT/07/IYBA/2013). The project was supported by Department of Biotechnology Membrane Structural Biology Programme Grant (BT/PR28833/BRB/10/1705/2018) to PG and funds from IISER Pune MB and SH. SH thanks Infosys Foundation (IISER-P/InfyFnd/ Trv/1) for fellowship.

## Author contributions

MB designed and performed all the experiments related to Fibril EGFP work, analyzed the data and wrote the manuscript. SH designed and performed all the experiments related to short constructs of Fibril and helped in initial standardization of Fibril EGFP work and wrote and reviewed the manuscript. RK helped with initial cloning of fluorescent constructs and purification of the short constructs. PG conceptualized, supervised and funded the project and wrote and reviewed the manuscript.

## Supplementary Figures

**Figure S1:**
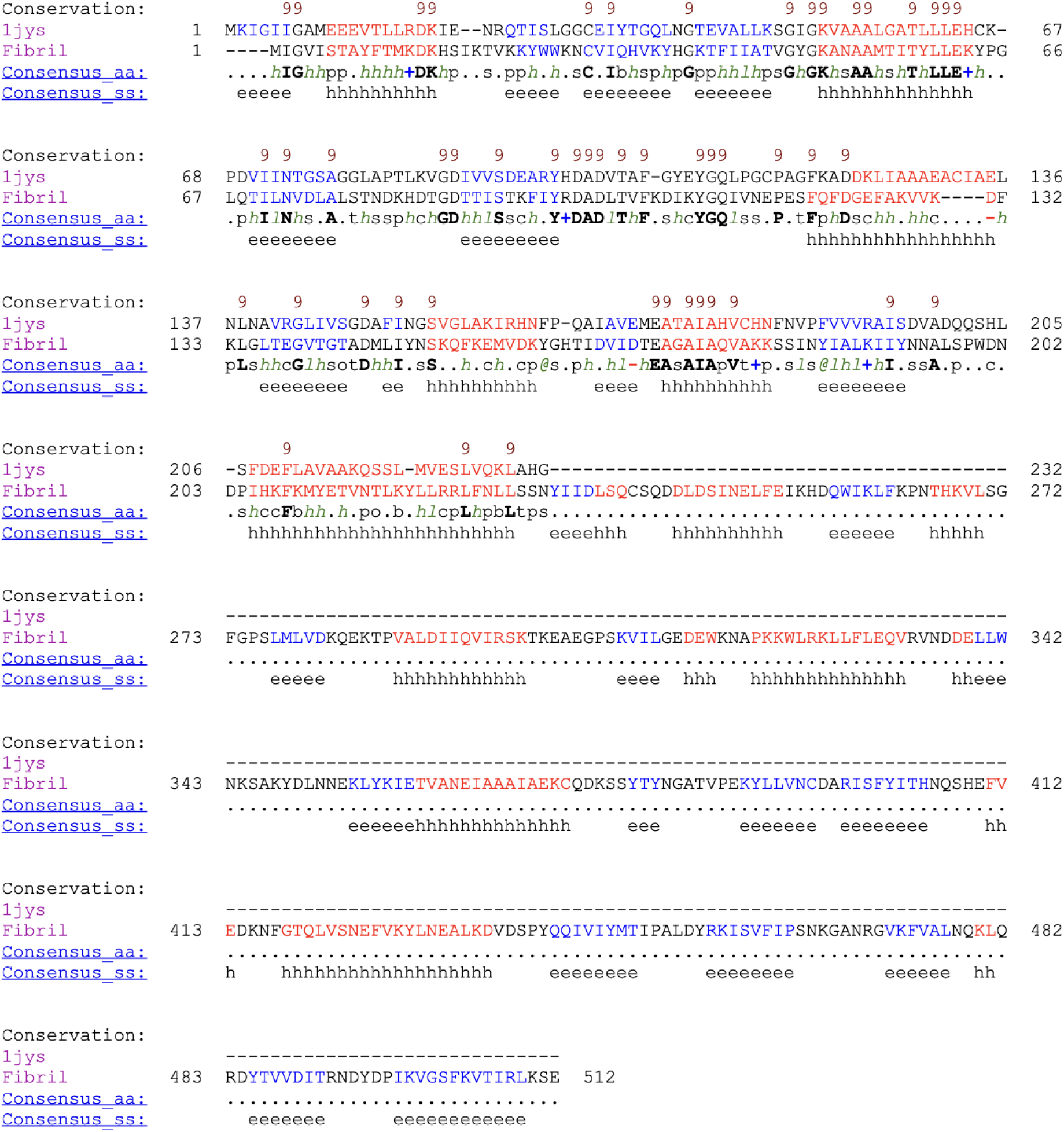
Prediction of Fibril domains. Sequence alignment of Fibril form *Spiroplasma citri* and 5’-methylthioadenosine/S-adenosylhomocysteine nucleosidase MTAN (PDB-1JYS) shows ∼24% sequence similarity. Alignment was done using Promals3D (http://prodata.swmed.edu/promals3d/promals3d.php).

**Figure S2:**
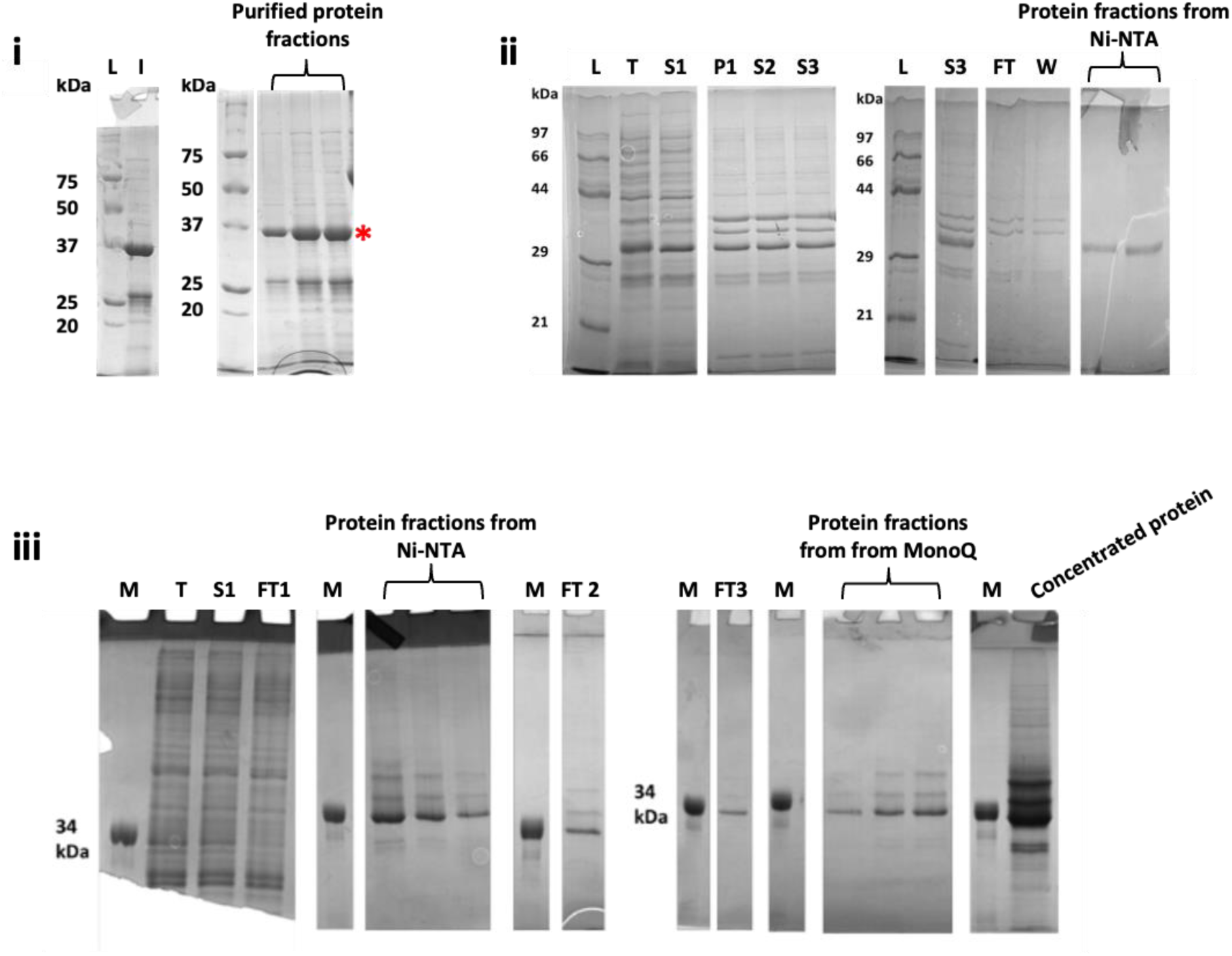
Purification of short constructs. **A. i.** The protein partially purified from pelleting assay of N-His_6_ Fibril(Δ305-512) was further purified by glycerol density gradient. **L**-Ladder, **I**-input obtained from pelleting assay of N-His_6_ Fibril(Δ305-512). The fractions of purified protein. **ii.** FibrilΔ(273-512)-His_6_ purification using Ni-NTA affinity chromatography. **L**-Protein marker; **T**-total lysate; **S1**-supernatant after spinning lysate at 11,000 xg; **P1**-Pellet obtained upon spinning protein as in lane 2 at 46,500 xg; **S2**-soluble fraction obtained upon solubilization of the pellet (lane 3) by treatment with SDS followed by spinning at 21,000 xg; **S3**-supernatant obtained by spinning sample from lane 4 at 159,000 xg; **FT & W**– flowthrough and wash (protein not bound to Ni-NTA matrix); proteins fractions bound and eluted using 250 mM and 500 mM imidazole containing buffer. **iii.** The protein FibrilΔ(273-512)-His_6_ was using Ni-NTA affinity chromatography. **L**-34 kDa protein; **T**-total lysate; **S1-** supernatant after spinning lysate at 46,500 xg; **FT1**-flowthrough from Ni-NTA column; protein fractions bound to Ni-NTA column and eluted using imidazole in the buffer; **FT2**-flowthrough from cation exchange (MonoS) column. The flowthrough of MonoS column was passed through anion exchange (MonoQ) column, bound protein eluted and concentrated. **L**-34 kDa protein, **FT3**-flowthrough of MonoQ column; fractions of protein bound to MonoQ column matrix and eluted with increasing salt concentration; Pooled and concentrated fractions from MonoQ column containing protein of interest.

**Figure S3:**
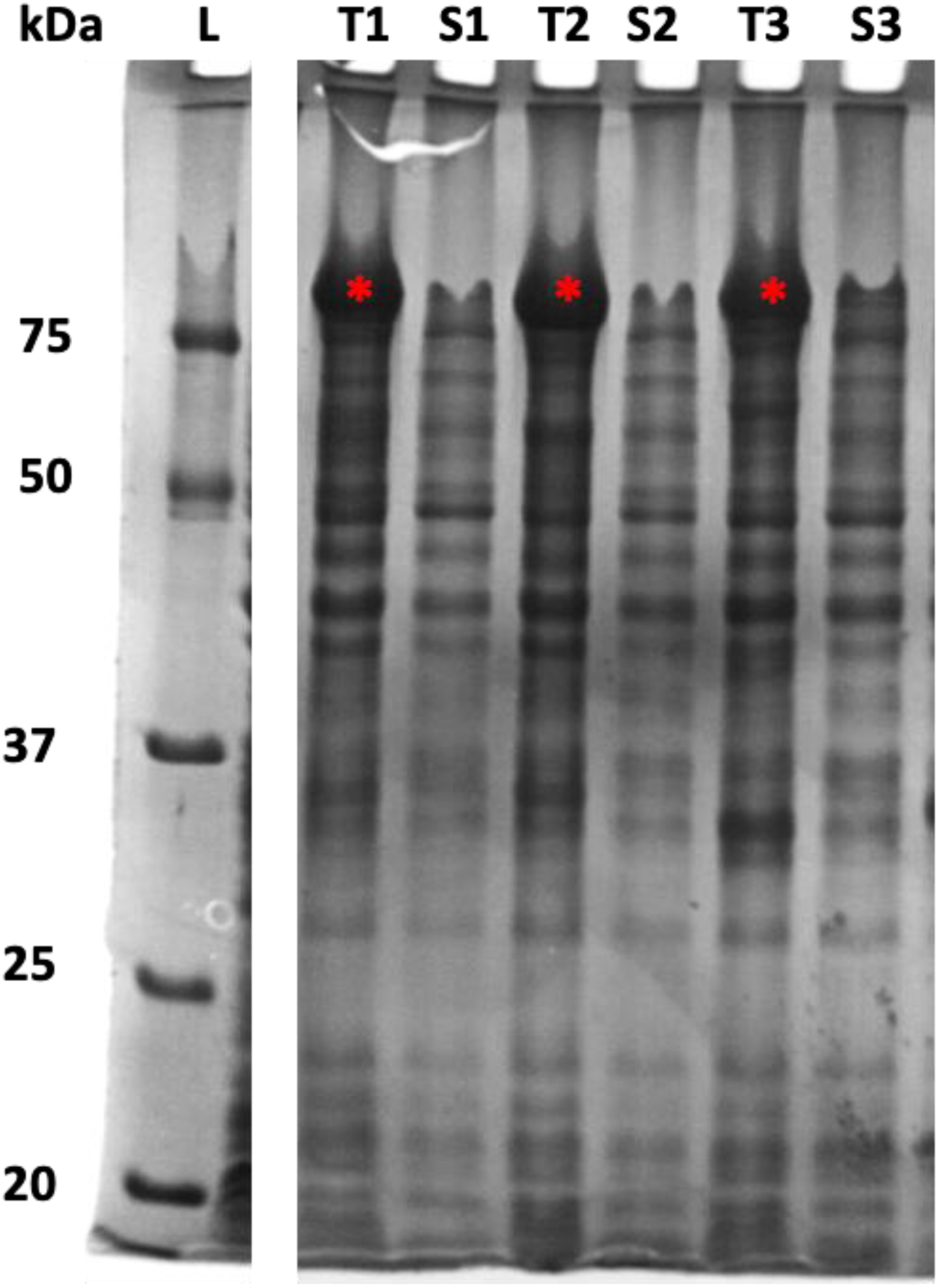
Representative gels of 3 Fibril-EGFP constructs. Fibril-EGFP fusion protein molecular weight is 86.6 kDa. Overexpression of 3 Fibril-EGFP constructs is shown in the gel. Lanes T1, T2 and T3 are induced total lysate samples. Lanes S1, S2 and S3 represents induced supernatant samples.

**Figure S4:**
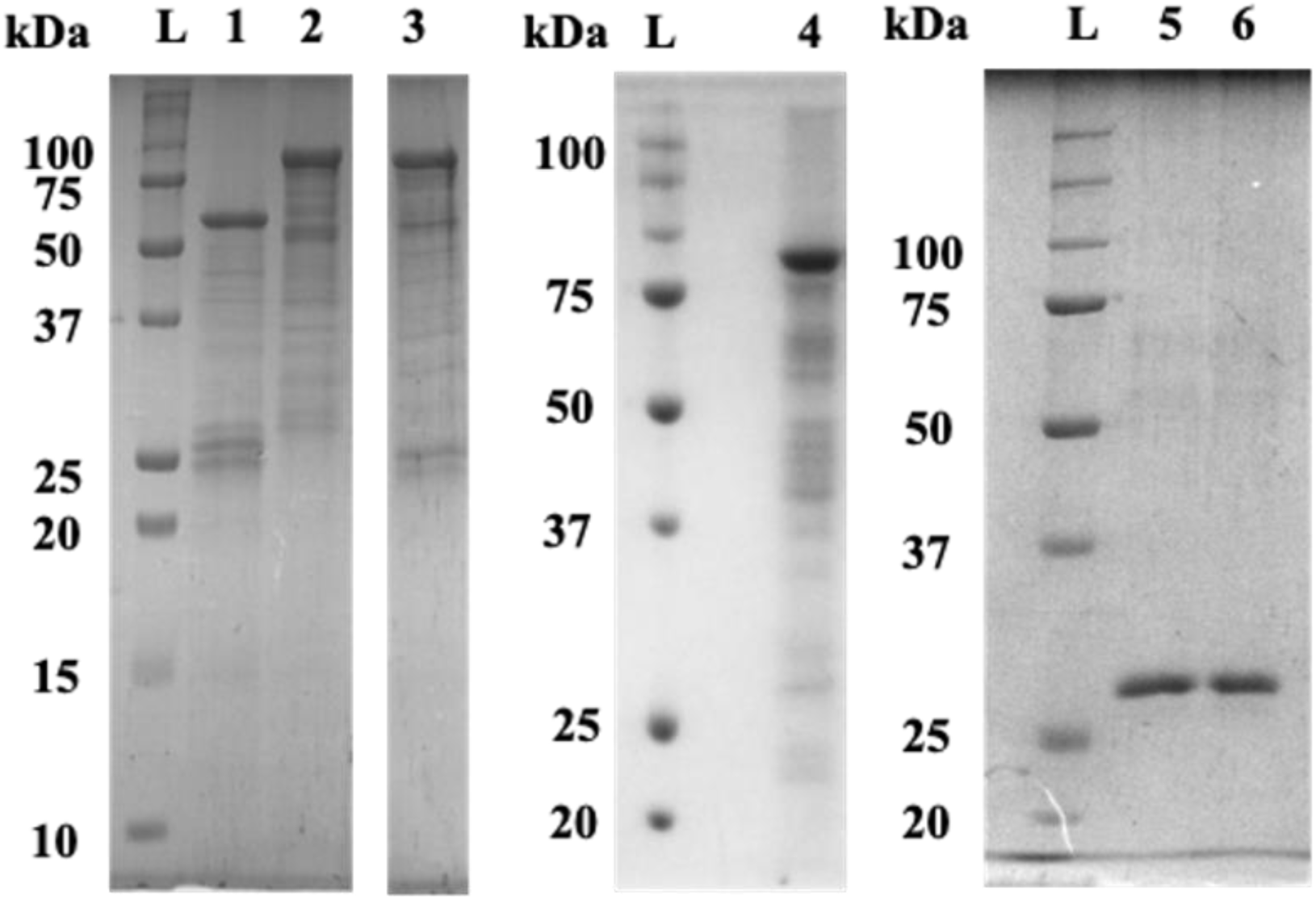
Concentration check of samples used for circular dichroism. Labels above the gels signify as follows, M-marker, 1-Fibril^WT^, 2-Fibril 27 EGFP, 3-Fibril 493 EGFP, 4-Fibril 183 EGFP, 5-EGFP with 1% SDS and 6-EGFP without 1% SDS. Samples used for CD were loaded on 10% SDS-PAGE gel. 5 μM of protein was taken for obtaining the spectra. For solubilization of Fibril, 1% SDS was added to all CD samples containing Fibril.

**Figure S5:**
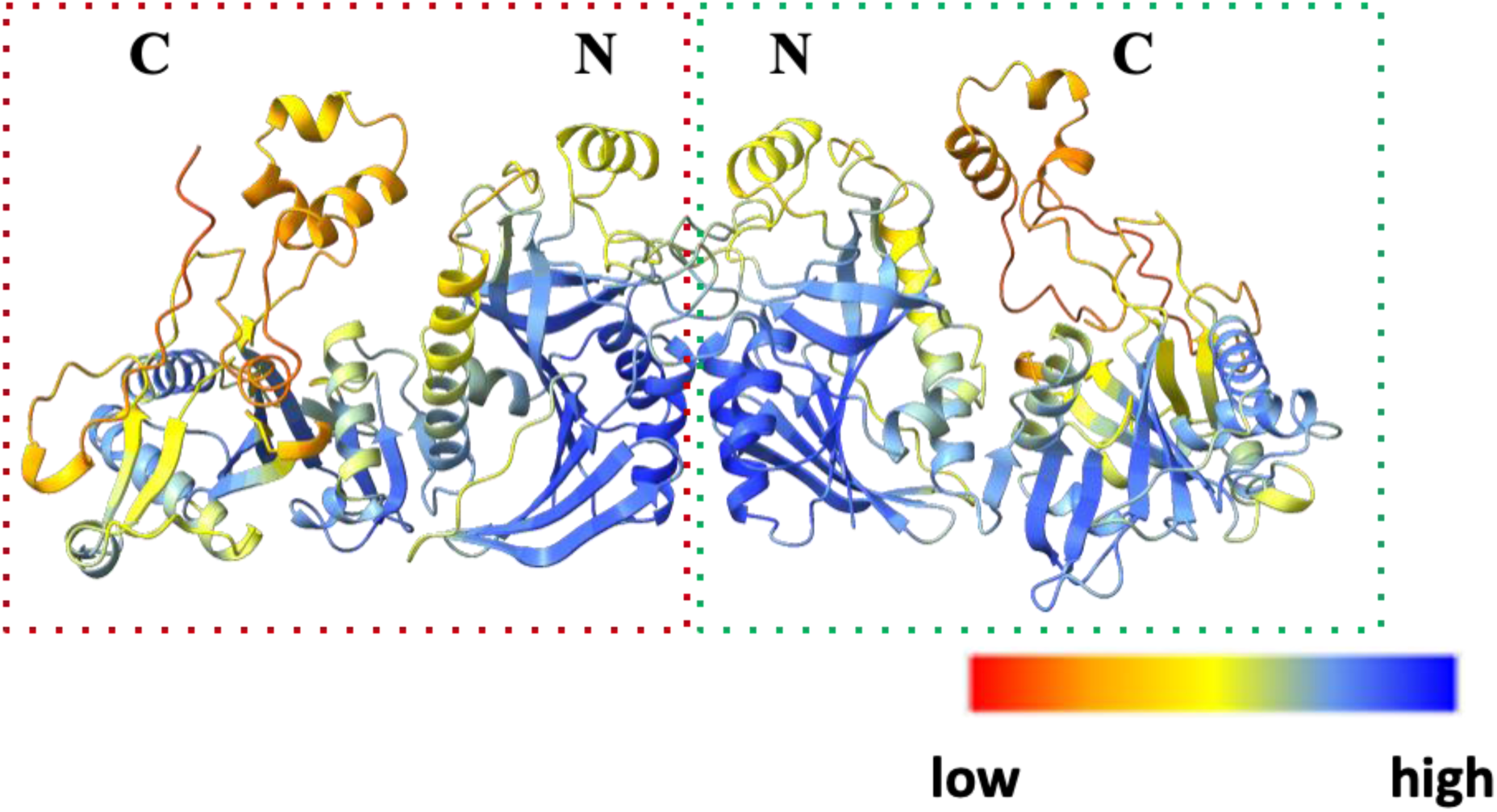
AlphaFold model for Fibril shows dimer interface as a part of N-terminal domain alone. Two monomers of Fibril are shown in red and green box each. The domains of each monomer has been denoted by N for N-terminal domain and C for C-terminal domain. Dimer model of Fibril predicts the dimer interface as a part of N-terminal domain alone. The dimer structure is colored according to the pLDDT score. The N-terminal domain shows high pLDDT score as compared to C-terminal domain.

**Table S1:**
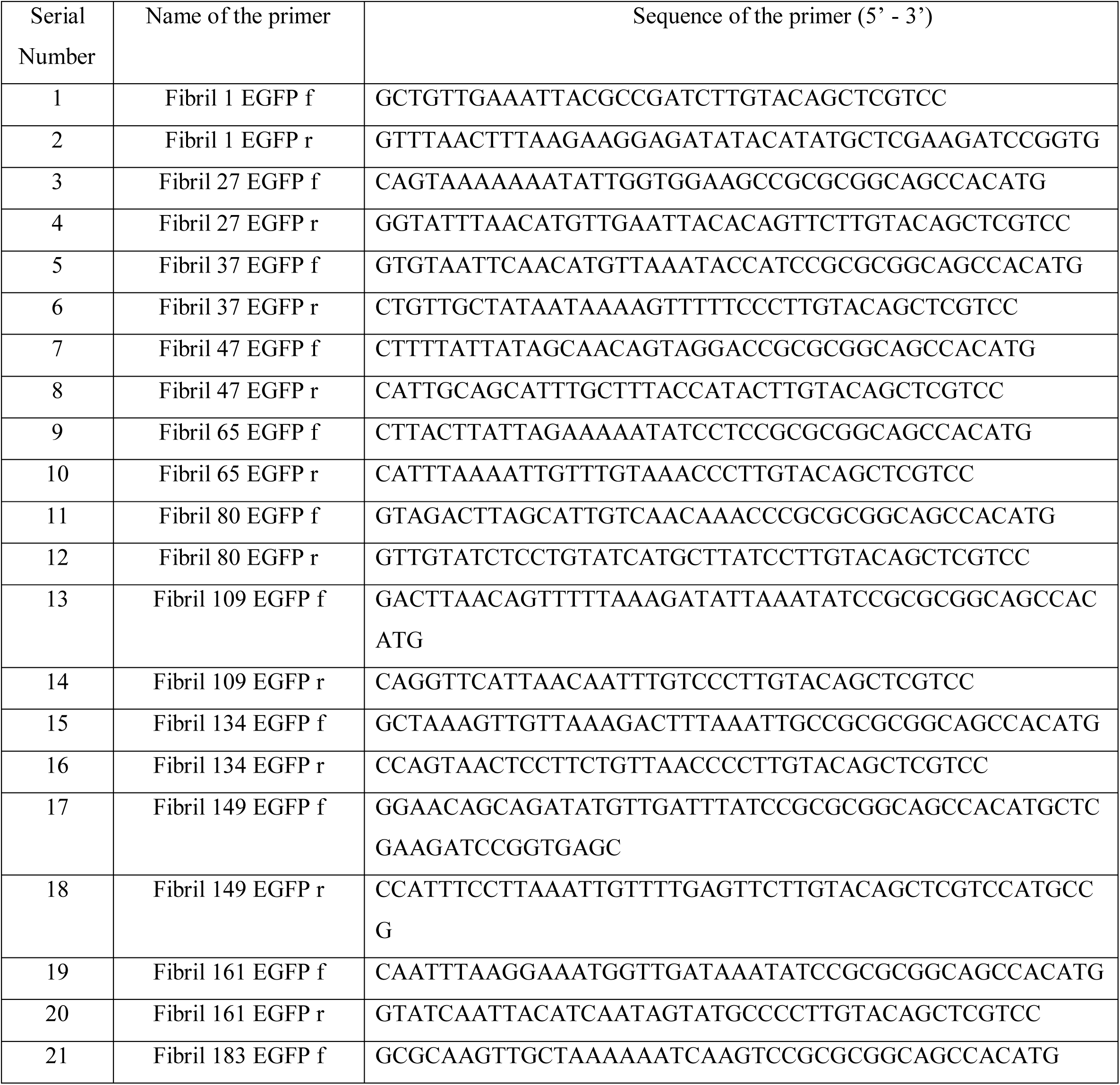
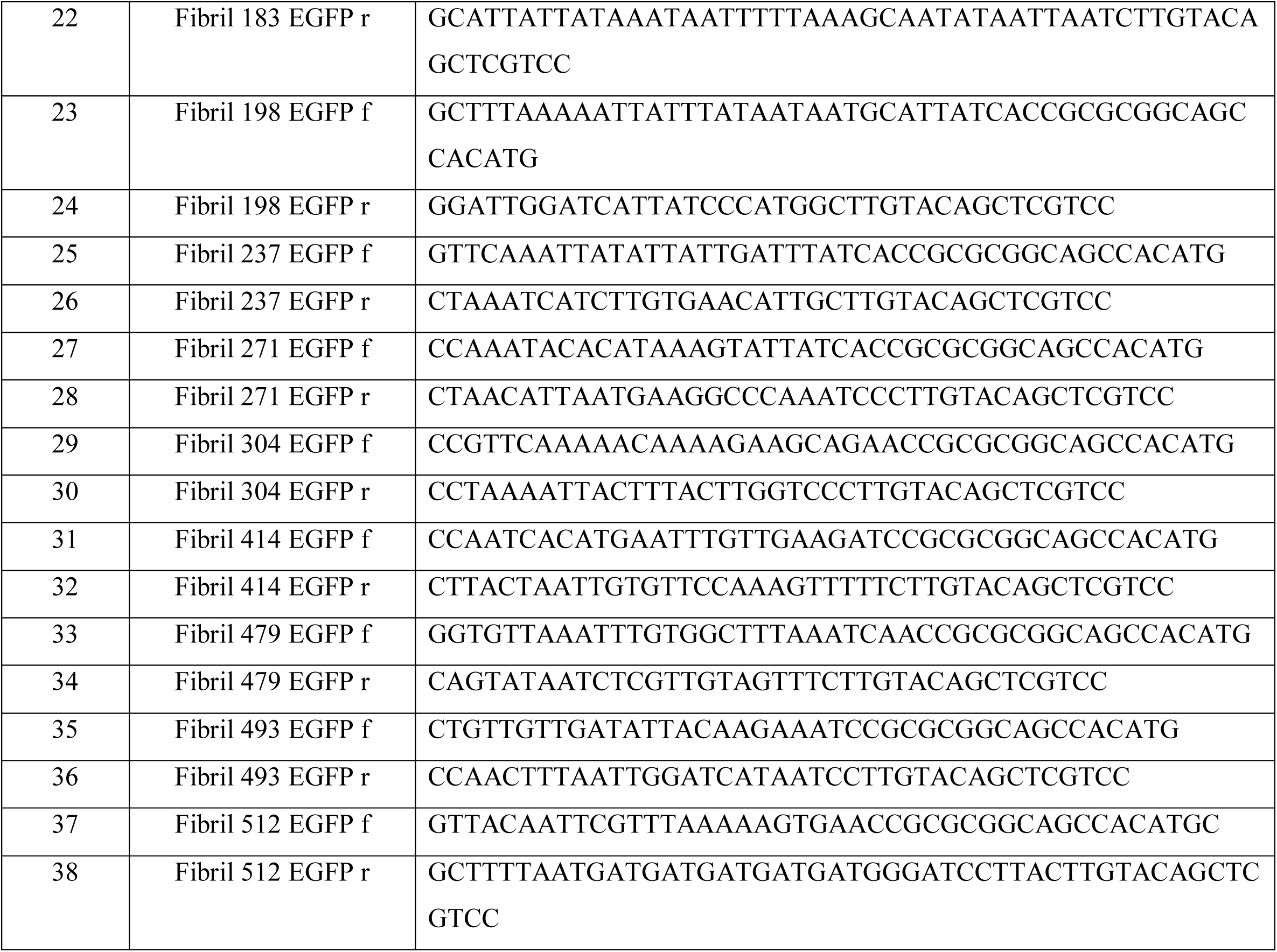
List of primers for Fibril-EGFP cloning.

## Notes

### Competing Interest Statement

The authors have declared no competing interest.

